# VEGFR-2 Phosphorylation at Y1054 or Y1214 is Necessary for Mechanically-Induced Angiogenesis

**DOI:** 10.64898/2026.08.21.746225

**Authors:** Bronte Miller Johnson, Tori McKinley, Thomas Nguyen, Eden Beasley-Duncan, Tanvi Giridhar, Mary Kathryn Sewell-Loftin

## Abstract

Anti-angiogenic cancer therapies attempt to withhold necessary nutrients and oxygen from growing tumors by targeting the major promoters of endothelial cell (EC) angiogenesis: vascular endothelial growth factor (VEGF) and VEGF receptor 2 (VEGFR-2). Unfortunately, these treatments are often insufficient, even when coupled with chemotherapies, and fail to significantly increase survival rates. These therapies focus on biochemical or ligand activation of VEGFR-2 and ignore mechanical cues which have been shown to promote angiogenesis. The tumor microenvironment (TME) is mechanically distinct compared to normal tissue, including increased matrix deformations or strains caused by cancer-associated fibroblasts (CAFs). Prior studies have shown that CAF-like strains activate VEGFR-2, which is not blocked by soluble inhibitors of the receptor. In this report, we detail the specific and independent roles of two tyrosine residues, Y1054 and Y1214, on mechanical activation of VEGFR-2 as related to strain-induced angiogenesis. Furthermore, we characterize CAF biochemical and mechanical signaling and demonstrate how ECs exhibit decreased vessel growth when co-cultured with CAFs and treated with a contractility inhibitor. Using non-phosphorylatable VEGFR-2 mutants, we reveal Y1054 and Y1214 are each necessary for EC angiogenesis, particularly in response to strain. Overall, this research highlights the need to study how mechanics in the TME promote vessel growth and thus tumor progression, which is important to consider when developing future anti-angiogenic therapies.

Graphical Abstract
Spectrum of angiogenic behaviors related to VEGFR-2 signaling initiated by either ligand, strain, or combinatorial cues. The ligand (red circle) activates phosphorylation at specific sites, while strain (black arrow, e) activates at different sites and prolongs activation. Phosphorylation represented by purple (P). Inhibition, such as clinically utilized bevacizumab, represented by white/green pill. These inhibitors are primarily suitable for blocking ligand-induced VEGFR-2 activation, and have little effect when strain in present. Bottom: representative figure of how much angiogenesis is stimulated by ligand and/or strain, with and without inhibitors.

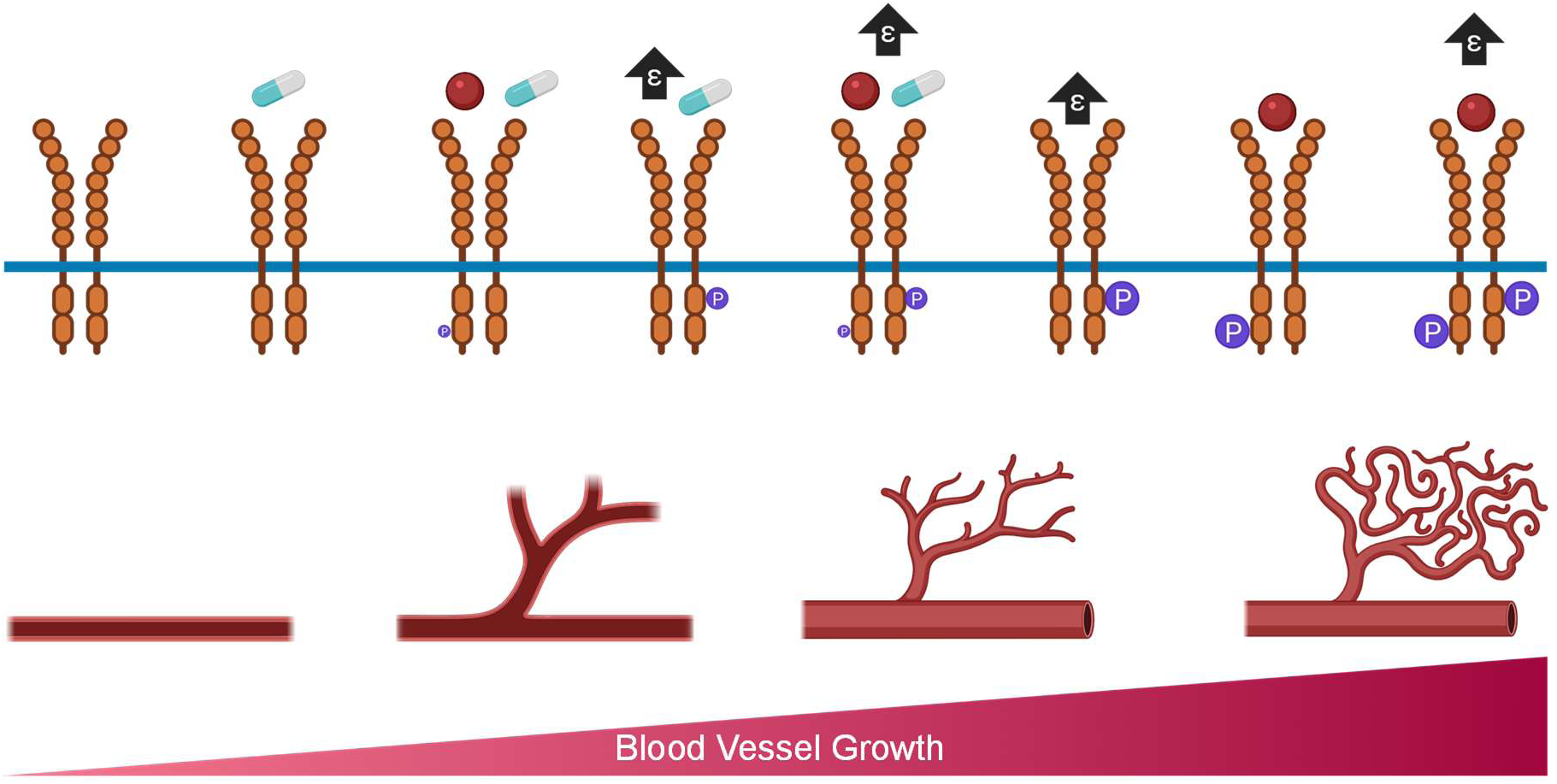

## Introduction

Blood vessel growth is an essential biological process required for development and wound-healing, but it is also necessary for tumor progression. Angiogenesis, the growth of new vessels from existing vasculature is initiated by numerous cues in a tissue including hypoxia inducible factors (HIFs), vascular endothelial growth factor (VEGF), and via beta-1 integrin signaling stimulated by interstitial fluid flow (Shirure et al., 2017, Hashimoto and Shibasaki, 2015). Angiogenesis is initiated when a quiescent endothelial cell is stimulated by one of these cues to become a tip cell, losing its cell-cell adhesions, becoming highly migratory, and initiating matrix remodeling (Miller and Sewell-Loftin, 2022). This process is different from vasculogenesis, which is blood vessel development without the pre-existing vasculature. While vasculogenesis is primarily associated with tissue development, it can also occur in some tumor microenvironments (TMEs) (Folberg et al., 2000). The fundamental biological processes of vasculogenesis and angiogenesis are designed to deliver enhanced nutrients and oxygen to tissues as they undergo either physiological or pathological remodeling. For tumor progression, when a tumor is small enough, diffusion is sufficient to supply oxygen and other nutrients. However, as the tumor becomes larger than this size threshold, it promotes angiogenesis from the surrounding healthy tissues to supply the needed nutrients (Carmeliet and Jain, 2000, Lugano et al., 2020). This angiogenic growth of blood vessels into the TME is considered a hallmark of tumor progression and, similar to many disease states, is widely dysregulated, contributing to worse clinical outcomes. Tumor and stromal cells begin to secrete VEGF, which binds to VEGF receptor 2 (VEGFR-2) on endothelial cells (ECs) and causes vessels to grow toward and vascularize the TME, thus allowing tumor progression (Roskoski, 2007). VEGF and VEGFR- 2 are the major promoters of angiogenesis, which is vessel growth off pre-existing vasculature. (Folberg et al., 2000) Notably, the VEGF/VEGFR-2 signaling axis is important in vasculogenesis as well. Cancer therapies often target angiogenesis to stop tumor growth by limiting nutrient supply, but these treatments often are insufficient by themselves and do not significantly increase survival rates in many patient populations (Ribatti, 2016, Ribatti, 2010). For example, bevacizumab, a monoclonal antibody which binds to VEGF and interferes with VEGFR-2 activation, was the first FDA-approved drug for cancer treatment in 2004 (Dey et al., 2015, van Beijnum et al., 2015). It has been approved to treat several cancers including lung, colorectal, and breast cancer; however, the approval to treat metastatic breast cancer was revoked in 2011 due to the inefficacy of the drug in this population (Sasich and Sukkari, 2012). While anti-VEGF/VEGFR-2 agents show great promise in preclinical models, there is less efficacy in specific patient populations, which discrepancy is not well understood. This study investigates how mechanical strains promote blood vessel growth via mechanoactivation of VEGFR-2 and subsequent responses to anti-VEGF/VEGFR-2 therapies.

Cancer therapies often overlook the role of mechanical forces, which in normal and tumorous tissues are well-known regulators of cellular behaviors. ECs experience several physical forces such as increased shear stress, interstitial fluid pressure, and stiffness (Yu et al., 2011, Roman and Pekkan, 2012, Ayad et al., 2019). In tumors, ECs also face increased mechanical strain due to the presence of cancer-associated fibroblasts (CAFs), highly contractile stromal cells that pull on surrounding matrix and cells (Zanotelli and Reinhart-King, 2018, Dessalles et al., 2021). Previous research has confirmed that CAFs compared to normal breast fibroblasts (NBFs) cause more extracellular matrix (ECM) deformation and promote more vessel growth when co-cultured with ECs (Sewell-Loftin et al., 2017, Koch et al., 2020, Bates et al., 2023, Alcoser et al., 2015). Mechanical strain has been shown to increase EC vasculogenesis and angiogenesis through the use of 3D microtissue models with fibrin gels (Johnson et al., 2023). Other physical forces affect VEGFR-2 activity. For example, one study observed that ECs grown on stiffer substrates displayed higher phosphorylation of VEGFR-2 Y1175 (LaValley et al., 2017b). ECs also respond to flow, with cells aligning perpendicular to the flow in high-VEGF conditions and vessel growth generally moving against flow direction (Vion et al., 2020, Shirure et al., 2017).

For activation and downstream signaling to occur, VEGFR-2 dimerizes on the cell surface upon VEGF binding to the extracellular domain (Koch and Claesson-Welsh, 2012). Phosphorylation occurs on the intracellular domain of VEGFR-2 at one or several different tyrosine residues and can subsequently cause VEGFR-2 internalization and degradation (Simons, 2012, Labrecque et al., 2003, Ewan et al., 2006). We have focused on VEGFR-2 phosphorylation sites Y1054/Y1059 and Y1214. Y1054/Y1059 are on the activation loop of VEGFR-2 and are necessary for full kinase activity (Kendall et al., 1999). Y1214 is linked to several signaling pathways, including cdc42, p38, and ERK (Lamalice et al., 2004, Testini et al., 2019). One study found that matrix-bound VEGF caused prolonged phosphorylation of Y1214 compared to soluble VEGF, suggesting that VEGFR-2 responds to mechanical cues (Chen et al., 2010).

Due to the several studies that demonstrate mechanoactivation of VEGFR-2, we decided to expound upon this topic and highlight the previously understudied effects of strain on angiogenesis in the TME. Better understanding how strain from CAFs affects VEGFR-2 signaling will provide insight into how to inhibit this pathway to develop more effective cancer treatments. This research characterizes the roles of Y1054/Y1059 and Y1214 and how these phosphorylation sites are necessary for vasculogenesis and angiogenesis, which are essential functions in tumor progression. Future anti-angiogenic therapies should consider these mechanical properties and target both the physical changes in the TME and the downstream signaling pathways that are activated by the altered mechanics. We hypothesized that mechanical activation of VEGFR-2 is not responsive to traditional inhibition strategies and explored the specific role of mechanically-induced phosphorylation events in vessel growth.

## Results

### Inhibition of Contractility Minimally Alters VEGF Secretion and Decreases Vasculogenesis

To begin to separate biochemical from biomechanical cues, we first measured secreted VEGF in CAFs treated with blebbistatin; our results show that there is a small but significant increase of VEGF measured in the media for CAFs treated with 20um blebbistatin compared to control or vehicle-treated groups (**Fig 1A**). To ensure that CAF-secreted VEGF was the primary biochemical driver, we analyzed conditioned media from 2D and 3D ECs. The results demonstrate that CAFs secrete roughly 1ng/mL VEGF, over 10-fold the amount of VEGF compared to any of the HUVEC samples **(Fig S1A)**. Next, we wanted to focus on the mechanical stimulation generated by CAFs, specifically their contractility and how to inhibit that contractility. CAFs in 3D microtissues containing tracer beads were treated with no treatment (NT), a DMSO vehicle control, or the contractility inhibitor blebbistatin at 20µM or 50µM for 24hr, then bead displacement was measured over 30min **(Fig 1B,C)**. Quantification of these displacements displays a significant decrease in matrix deformation generated by the CAFs when treated 20µM or 50µM blebbistatin compared to relevant controls. In vasculogenesis studies, microtissues containing CAFs and HUVEC-TERT2 cells were cultured for 3d in standard media before addition of vehicle or 20um blebbistatin for 4d; these studies demonstrated significant decreases in blood vessel growth per area for blebbistatin-treated samples (**Fig 1D,E**). CAFs and HUVECs grown in 3D and treated with either 20uM or 50uM of blebbistatin did not show significantly higher cell death compared to cells treated with a vehicle control, ruling out cell death as a reason for decreased vasculature with blebbistatin treatment **(Fig S1B,C)**.

**Fig. 1.**
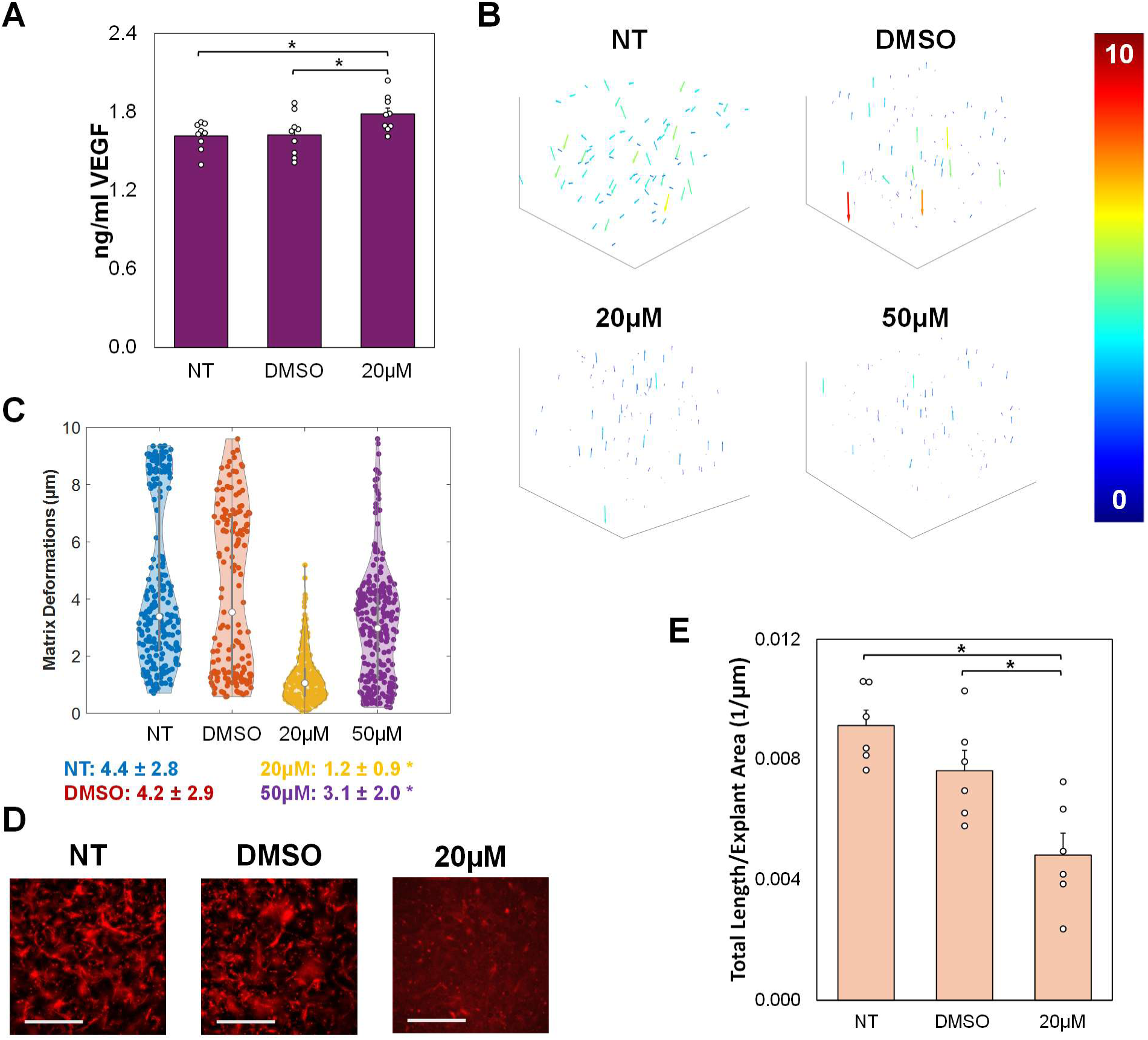
CAF Ligand-Based and Mechanical Signaling. **(A)** Conditioned media was harvested from 3D CAFs in fibrin gels after 48hr of treatment with no treatment (NT), a vehicle control (DMSO), or 20µM blebbistatin for analysis via VEGF ELISA. Data shown as average + SEM, n=9. *p<0.05. **(B)** A matrix deformation assay was performed with CAFs with no treatment (NT), vehicle control (DMSO), 20µM blebbistatin, or 50µM blebbistatin. MATLAB figures show matrix deformations represented by arrows. Colors correspond to magnitude (0-10µm). **(C)** Data from images in (b) presented in violin plots. Average values are represented by white dots; average ± SD shown as insets below figure; n>30 bead tracings in 2- 3 biological replicates. *p<0.05 compared to NT and DMSO. **(D)** Representative images of 3D fibrin rings containing HUVEC TERT2s co-cultured with CAFs treated with NT, DMSO, or 20µM blebbistatin. Gels were stained for CD31 (red). Scale bar=500µm. **(E)** Graph of total vessel length divided by explant area, n=6. Graphs show average + SEM. Data in (A) and (E) were compared with ANOVA, followed by post hoc Tukey HSD tests. Data for C) was compared with Kruskal-Wallis test, followed by post hoc Dunn’s tests.

### VEGFR-2 Y1054 and Y1214 Mutations Blunt EC Strain Response

Previously, our lab determined that strain is sufficient to induce VEGFR-2 activation and phosphorylation at Y1054/Y1059 or Y1214 independent of VEGF chemical signaling (Johnson et al., 2023). We next characterized two CRISPR/Cas9 mutant HUVEC TERT2 cell lines which replaced either Y1054 (Y1054F) or Y1214 (Y1214F) with a phenylalanine, which cannot be phosphorylated, using a Cas9 control line that contained no genetic modifications **(Fig 2A)**. Results show that 25ng/mL exogenous VEGF does not alter total VEGFR-2 expression in Cas9 or Y1054F (**Fig 2B**). However, 15min treatment with exogenous VEGF increases pY1054/Y1059 relative to total VEGFR-2 for Cas9 samples (**Fig 2C**). Our data show decreases in pY1214 levels normalized to VEGFR-2 in both Cas9 and Y1054F samples with VEGF treatment (**Fig 2D**). There are no significant differences observed in pY1054/Y1059 or pY1214 levels for Y1054F samples when normalized to β-actin, which may be due to phosphorylation still occurring at Y1059 (**Fig S2A,B**).

**Fig. 2.**
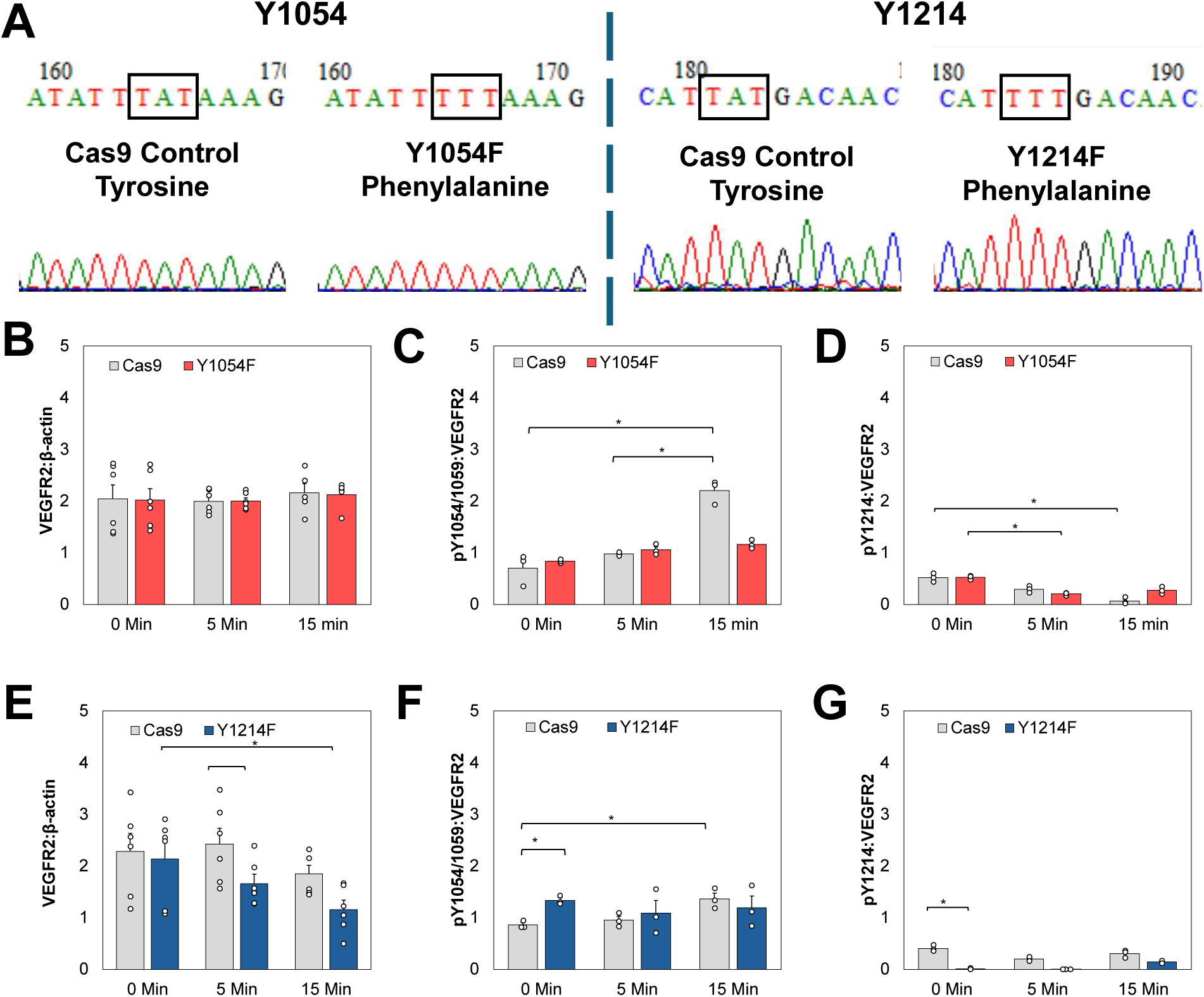
Y1054 and Y1214 Are Necessary for Activation of VEGR-2 through VEGF. (**A**) HUVEC TERT2 CRISPR/Cas9 mutant cell lines were developed changing either tyrosine 1054 or 1214 into a phenylalanine (Y1054F and Y1214F), which cannot be phosphorylated. Cas9 control lines were also utilized which have the Cas9 enzyme but no genetic mutation. Black boxes show the 1054 and 1214 codons which were mutated. (**B-D**) Y1054F and Cas9 lines were cultured on tissue culture plastic and then treated with 0, 5, or 15min 25ng/mL VEGF in EGM-2. Western blot results for (B) VEGFR-2, (C) pY1054/1059, and (D) pY1214 normalized to β-actin loading controls or total VEGFR-2. (**E-G**) Western blot results for VEGFR-2, pY1054/Y1059, or pY1214 for Y1214F samples normalized to β-actin loading controls or total VEGFR-2. Data shown as average + SEM, n=3-6;*p<0.05. All groups were compared with Kruskal-Wallis test, followed by post hoc Dunn’s test. Corresponding figures for pVEGFR-2 normalized to β-actin can be found in **Fig S2A-D**; full membrane images are available in Additional File 2.

Results show significant decreases in total VEGFR-2 levels for Y1214F samples compared to Cas9 cells after 5min of VEGF stimulation (**Fig 2E**). There are no significant changes in pVEGFR-2 relative to total VEGFR-2 expression at either location for Y1214F samples regardless of VEGF treatment (**Fig 2F**). When normalized to β-actin, pY1054/Y1059 levels significantly decrease over time in Y1214F cells (**Fig S2C**). Furthermore, pY1214 expression is significantly lower in Y1214F than in Cas9 cells when both are normalized to total VEGFR-2 and β-actin, confirming that the mutation successfully inhibited phosphorylation at the Y1214 residue **(Fig 2G, S2D)**.

Y1054F and Y1214F samples were also treated for 15min with VEGF (25ng/mL), Strain (ε), or VEGF and Strain (VEGF+ε); no treatment (NT) samples had 0ng/mL VEGF and were cultured on Flexcell plates as controls. In samples treated with Strain only, total VEGFR-2 levels are higher in Y1054F cells compared to Cas9 samples **(Fig 3A)**. There were no significant changes observed in pY1054/Y1059 levels for any of the treatment groups when comparing Cas9 and Y1054F samples (**Fig 3B**). However, when samples are normalized to β-actin, the VEGF plus Strain group demonstrates significantly decreased levels of pY1054/Y1059 in comparison to the Strain only group in Y1054F samples (**Fig S2E**). For all treatment groups, the Y1054F samples show lower pY1214 compared to the equivalent Cas9 samples (**Fig 3C**). When normalized to β-actin, pY1214 levels are lower in the Cas9 samples when compared to the Y1054F for all treatment groups (**Fig S2F**). Total VEGFR-2 levels are unchanged in Cas9 control and Y1214F samples regardless of treatment (**Fig 3D**). In Y1214F samples, no significant differences were seen in pY1054/1059 or pY1214 levels when normalized to total VEGFR-2 or β-actin across all treatment groups (**Fig 3E, F, S2G**).

**Fig. 3.**
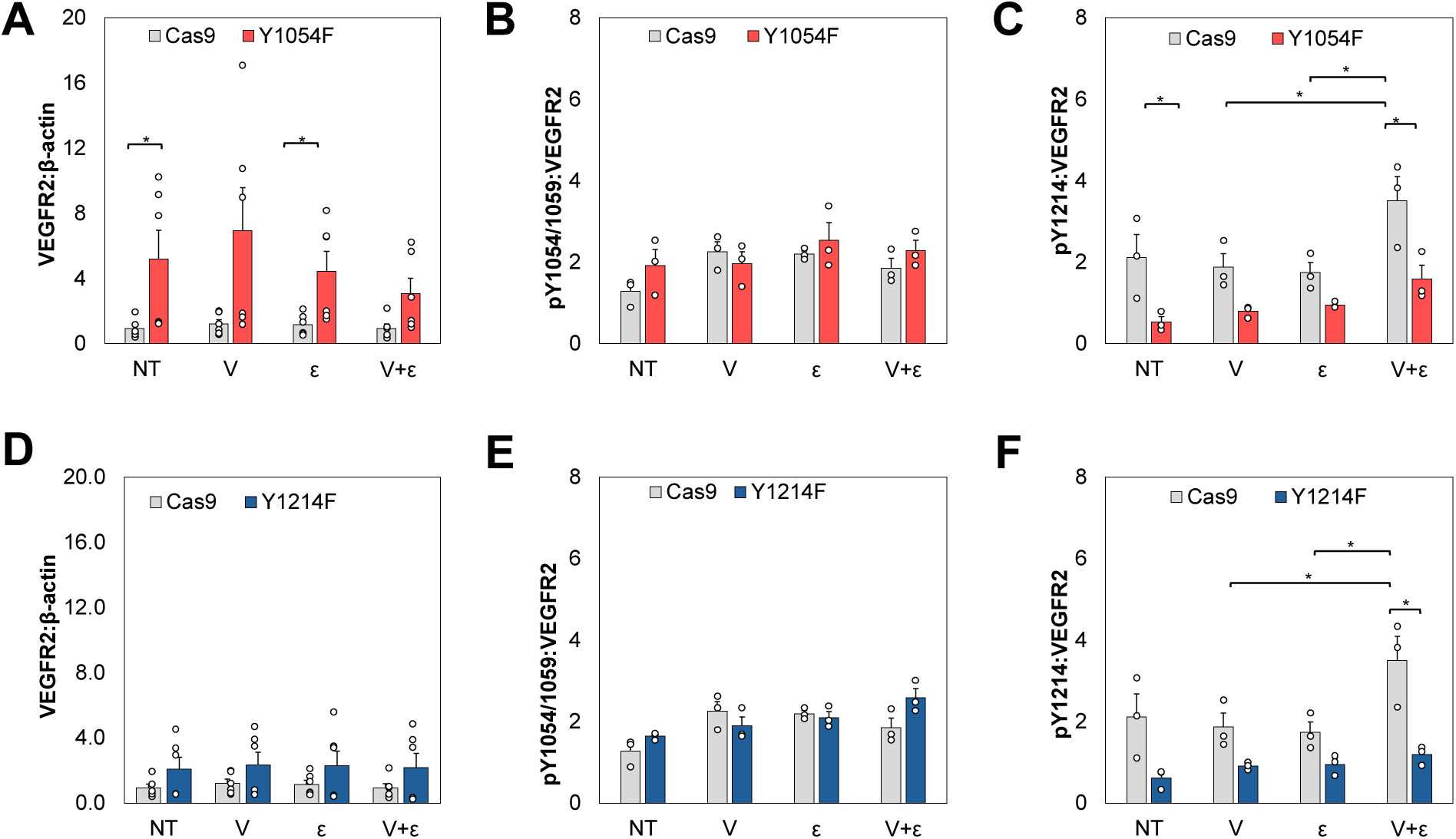
Y1054 and Y1214 Are Needed for VEGFR-2 Strain Response. (**A-C**) Y1054F and Cas9 lines were cultured on collagen-coated Flexcell plates and then treated with 15min 25ng/mL VEGF in EGM-2 (V), strain (s), combination VEGF and strain (V+s); control samples received no treatment (NT). Western blot results for (B) VEGFR-2, (C) pY1054/1059, and (D) pY1214 normalized to β-actin loading controls or total VEGFR-2. (**D-F**) Western blot results for VEGFR-2, pY1054/Y1059, or pY1214 for Y1214F samples normalized to β-actin loading controls or total VEGFR-2.. Data shown as average + SEM, n=3- 6;*p<0.05. All groups were compared with Kruskal-Wallis test, followed by post hoc Dunn’s test. Corresponding figures for pVEGFR-2 normalized to β-actin can be found in **Fig S2E-H**; full membrane images are available in Additional File 2.

### Loss of Y1054 or Y1214 Decreases Vasculogenesis

Y1054F and Y1214F mutant lines along with the Cas9 control line were co-cultured with NHLFs in our 3D fibrin microtissue model for 7d to characterize the effects that these mutations have on vasculogenesis. NHLFs were selected as a non-cancer stromal cell due to their highly pro-vasculogenic behaviors (Bala et al., 2024). The average length of vessels in the Cas9 samples is significantly higher than either of the mutant lines, confirming that phosphorylation at Y1054 or Y1214 is necessary for normal EC cell vasculogenesis **(Fig 4A,B)**. Samples with mutant VEGFR-2 were unable to promote vessel growth in response to mechanical stimulation from magnetic beads **(Fig 4C,D)**. Overall, this 3D microtissue model demonstrates that phosphorylation of VEGFR-2 at Y1054 or Y1214 is essential for blood vessel growth, and these sites are responsible for promoting vasculogenesis in response to mechanical strain.

**Fig 4.**
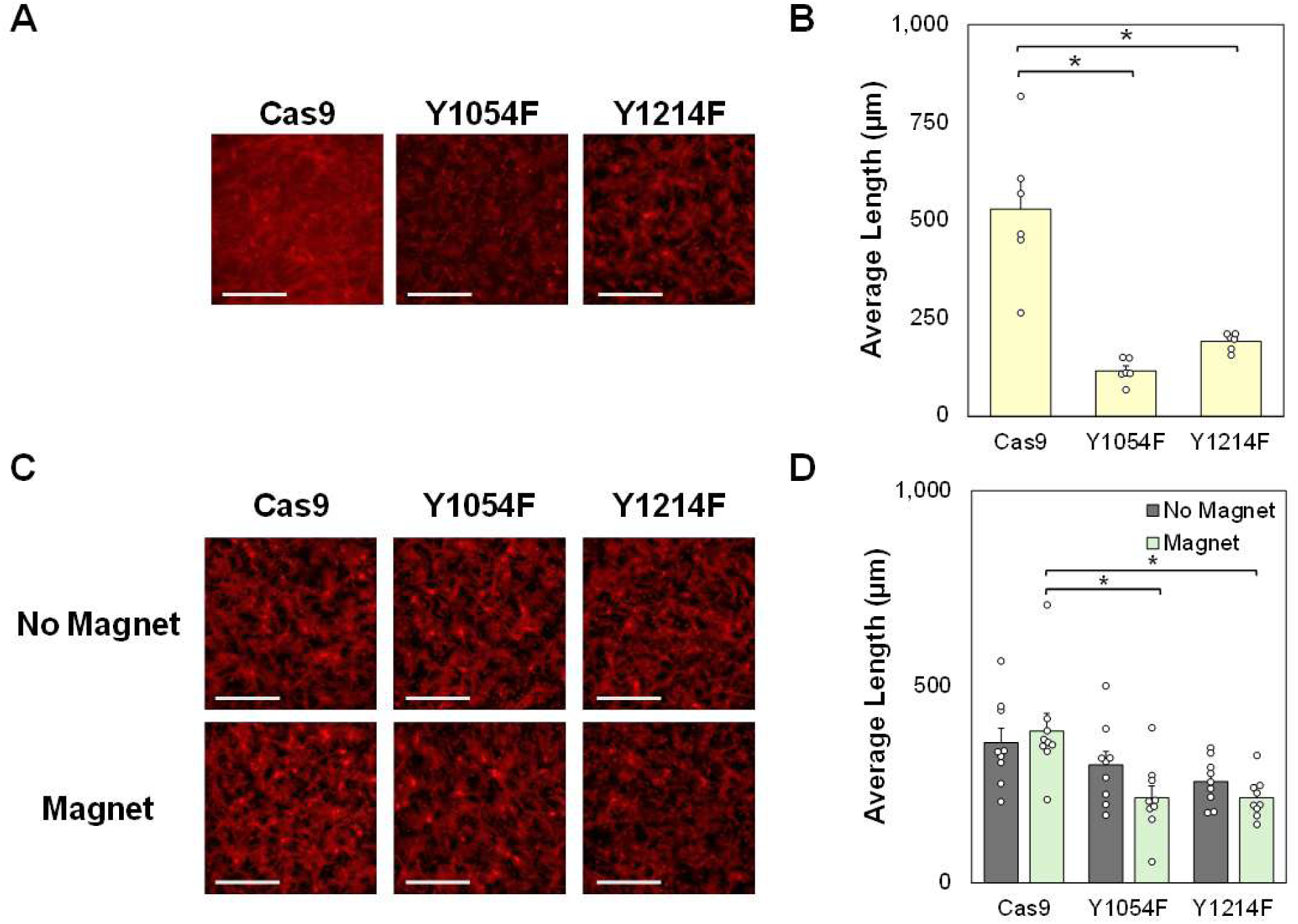
Phosphorylation at Y1054 or Y1214 Is Necessary for EC Vasculogenesis. **(A)** Representative images of HUVEC TERT2 Cas9, Y1054F, and Y1214F cells co-cultured with NHLFs in 3D fibrin rings for 7d then fixed and stained for CD31 (Red). **(B)** Quantification of average blood vessel length from studies shown in (A). **(C)** Representative images of HUVEC TERT2 Cas9, Y1054F, or Y1214F cells cultured for 7d with magnetic microbeads in 3D fibrin rings. Samples denoted as “Magnet” were grown above a rotating magnet, creating microstrains. **(D)** Quantification of average blood vessel length from studies shown in (C). Scale bars=500µm. Data shown as average + SEM; n = 6-9; *p<0.05. Data in (B) were compared with ANOVA, followed by post hoc Tukey HSD tests. Data in (D) were compared with Kruskal-Wallis test, followed by post hoc Dunn’s tests.

### Bevacizumab Fails to Block Mechanically-Stimulated Vessel Growth

VEGF ELISAs determined the bevacizumab concentration required to inhibit soluble VEGF in vessel assays. Treatment with increasing bevacizumab concentrations caused a dose-dependent decrease in VEGF levels in CAF and CAF+HUVEC TERT2 samples (**Fig 5A, S3A, S4A, Table S2**). A concentration of 25µg/mL of bevacizumab (Bev) was utilized in 3D vasculogenesis models with and without mechanical stimulation. After 7d in culture, results show that Y1054F samples generate shorter vessels compared to Cas9 controls (**Fig 5B, S3B,C, S4C, Table S2**). However, there are no significant differences in average vessel length for Y1214F samples when compared to Cas9 samples. In HUVEC TERT2 monocultures, treatment with Bev leads to a decrease in average vessel length (**Fig 5C**). Notably, vessel growth is rescued for samples that receive mechanical stimulation from magnetic beads even when treated with bevacizumab (**Fig 5C**).

**Fig. 5.**
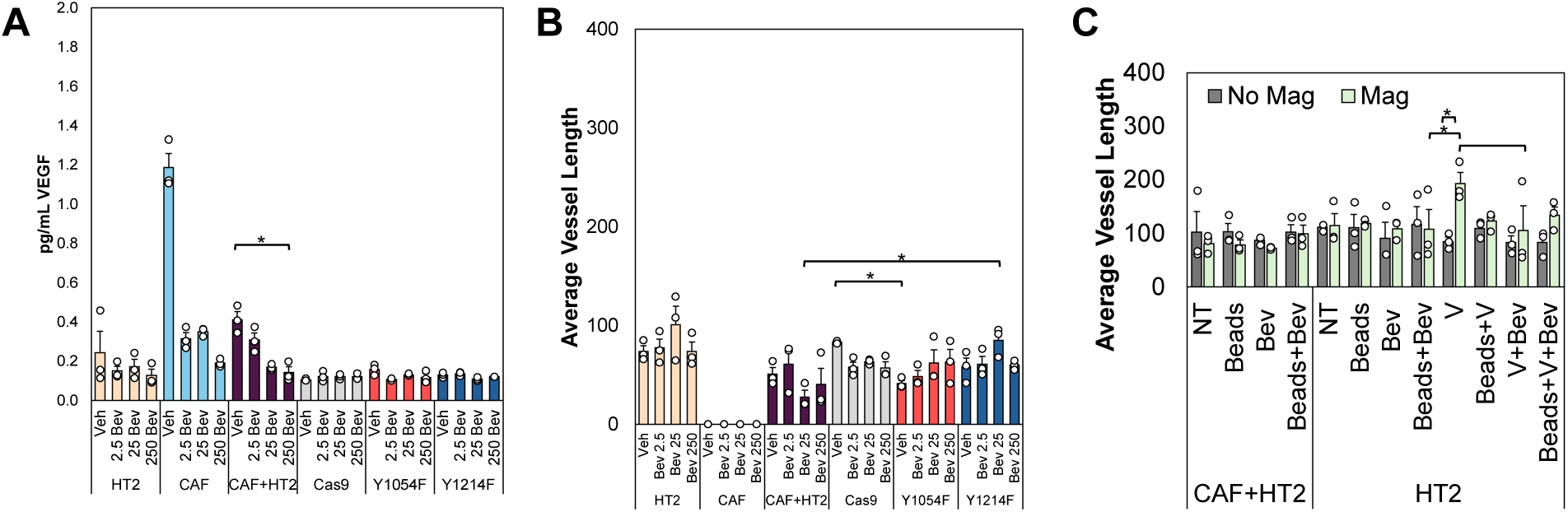
Bevacizumab Inhibits Endogenous VEGF Cues and Partially Blunts Vasculogenesis. (**A**) A VEGF ELISA was performed on conditioned media from HUVEC TERT2s, CAFs, HUVEC TERT2 Cas9, Y1054F, Y1214, or co-cultures of HUVEC TERT2 and CAF cells cultured in 3D fibrin rings. Full statistical data shown in Supplemental Table 1. (**B**) Average vessel length was quantified for 3D microtissues after 7d culture. All samples are significantly higher than CAFs, not shown for clarity of figure (**SI Table 2**). (**C**) HUVEC TERT2 and HUVEC TERT2 and CAF co-cultures in 3D fibrin disks were treated with no treatment (NT), magnetic microbeads (Beads), 25µg/mL Bevacizumab (Bev), 25 ng/mL VEGF (V), or combinations. Samples denoted as “Magnet” were grown on an external rotating magnet, creating microstrains. Average vessel length was quantified for studies with magnetic beads as mechanical stimulation, n=3. Data shown as average + SEM; n= 3;. *p<0.05. All groups were compared with Kruskal-Wallis test, followed by post hoc Dunn’s test. Note: HUVEC TERT2 abbreviated as HT2 in figures for brevity.

### Y1054 or Y1214 Necessary for Mechanically-Induced Angiogenesis

To determine the role of Y1054 and Y1214 in angiogenesis in these microfluidic devices, we co-cultured Cas9, Y1054F, or Y1214F cell lines with NHLFs in the center chambers of a three chambered microfluidic model previously developed (Sewell-Loftin et al., 2020). In one study, we loaded the side chambers with either NBFs or CAFs. As expected, the center chambers with the Cas9 cells display the highest vasculogenesis compared to Y1054F and particularly Y1214F cells **(Fig 6A,B)**. Our data also show that Cas9 ECs undergo the angiogenesis into both the NBF and CAF side chambers while the Y1054F and Y1214F samples show virtually no angiogenic growth into side chambers **(Fig 6C)**. To determine if mechanical stimulation alone could promote angiogenesis in the mutant lines, the magnetic bead assay was employed. Cas9 samples were able to undergo vasculogenesis in the center chamber and angiogenesis into side chambers with mechanical stimulation (**Fig 6D-F**). Both the Y1054F and Y1214F mutant lines demonstrate somewhat increased angiogenesis into side chambers with magnetic beads generating mechanical stimulation **(Fig 6F)**. Together, these results show that each tyrosine residue is able to promote some vascularization although this is decreased versus Cas9 control samples.

**Fig 6.**
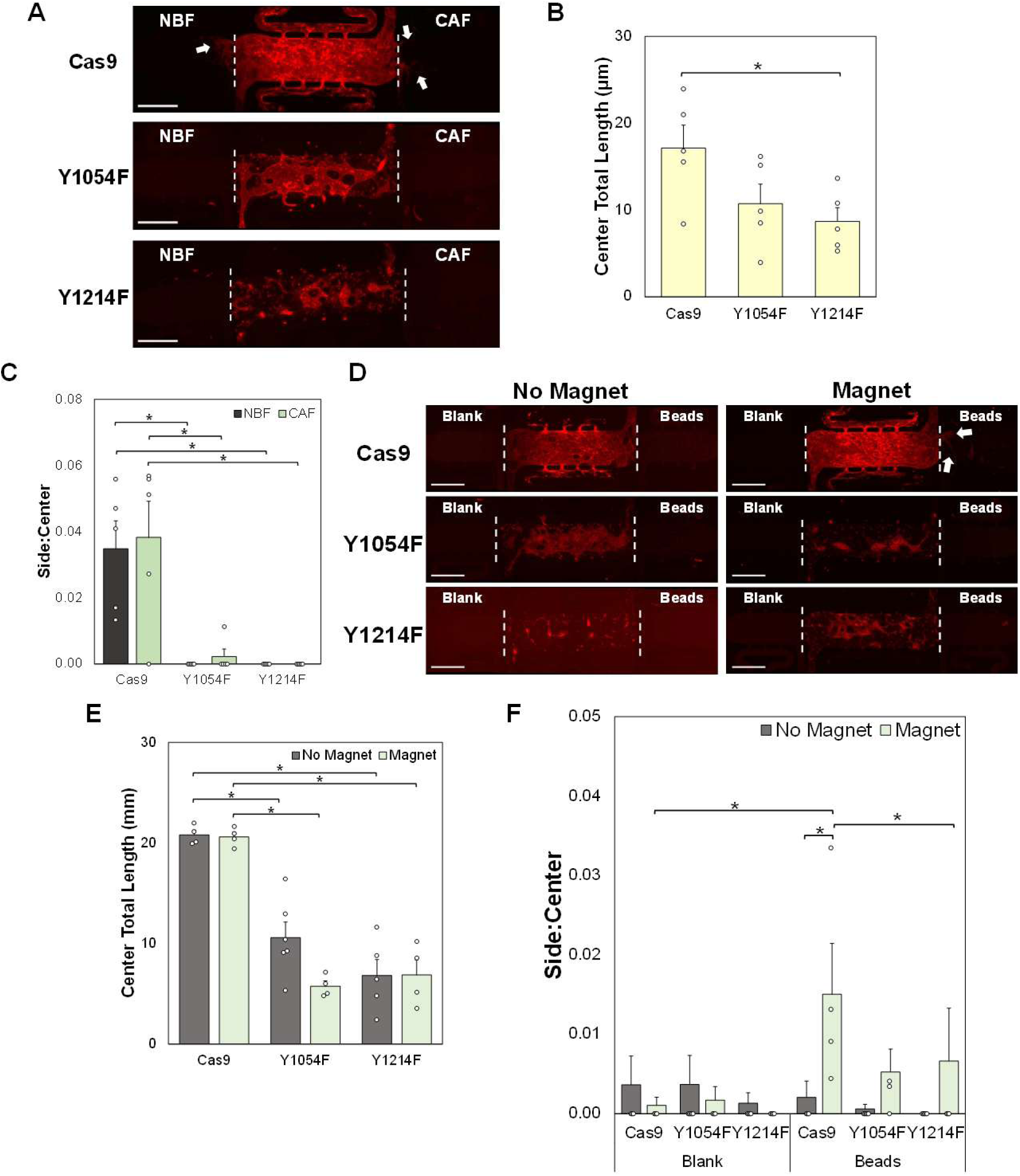
Phosphorylation at Y1054 or Y1214 Regulates Strain-Induced Angiogenesis. **(A)** Representative images of microfluidic devices were seeded with NHLFs and either HUVEC TERT2 Cas9, Y1054F, or Y1214F cells in the center chamber. For full device description, please see reference 38. The side chambers were loaded with NBF or CAFs. **(B)** Quantification of blood vessel growth in microfluidic models. Total vessel length in side chambers was normalized to total vessel length in the center chamber. **(C)** Representative images of microfluidic devices seeded with NHLFs and either HUVEC TERT2 Cas9, Y1054F, or Y1214F cells in the center chamber. Side chambers were loaded with a blank fibrin gel or magnetic microbeads. Samples noted “Magnet” were devices were grown above a rotating magnet, creating microstrains. **(D)** Representative images of microfluidic devices; center chambers were seeded with NHLFs and the mutant VEGFR-2 lines as noted. Insets represent side chamber loading conditions. Devices labeled as “Magnet” were cultured over a rotating magnetic field to generate microstrains. (**E**) Quantification of blood vessel growth in microfluidic center chambers, representing vasculogenesis. (**F**) Quantification of total vessel length in side chambers was normalized to total vessel length in the center chamber. Data shown as average + SEM, n=4-6; *p<0.05. Scale bars = 500µm. Data were compared with Kruskal-Wallis test, followed by post hoc Dunn’s tests.

## Discussion

This project highlights the important role of mechanical strain caused by CAFs in the TME, in promoting angiogenesis. Moreover, our results show that inhibiting only biochemical drivers of angiogenesis via bevacizumab may not be sufficient to block tumor-associated angiogenesis. Current anti-angiogenic therapeutic strategies mimic bevacizumab, by blocking binding of the ligand to the receptor, or attempt to interrupt the receptor tyrosine kinase activity of VEGFR-2. CAFs have been shown to both produce VEGF and generate microstrains in the ECM, and these functions likely interact to drive increased vessel growth observed in ECs co-cultured with CAFs **(Fig S1A)** (Sewell-Loftin et al., 2017). Therefore, biochemical and biomechanical cues work in concert to promote VEGFR-2 activation and subsequent blood vessel growth. Our studies demonstrated that CAFs secrete significantly higher levels of VEGF compared to ECs and that blocking cytoskeletal contractility with blebbistatin actually increases VEGF levels but decreases vessel growth (**Fig 1A, D, S1**). Results show that blebbistatin was not highly cytotoxic, so that observations about decreased contractility or vessel growth are not due solely to off-target effects that cause cell death (**Fig S1C**). These results showcase that mechanics within the TME should be considered when attempting to halt angiogenesis and tumor progression. Contractility inhibitors, along with VEGF or VEGFR-2 inhibitors, may be considered when developing future anti-angiogenic therapies, as it will aid in normalizing the TME and reducing mechanoactivation of VEGFR-2.

We previously studied the effects of strain on VEGFR-2 phosphorylation at Y1054/Y1059 and Y1214, finding that strain without any exogeneous VEGF is sufficient to cause phosphorylation and angiogenesis (Johnson et al., 2023). Other research has shown that Y1054/Y1059 is necessary for full kinase activity of VEGFR-2 (Kendall et al., 1999). Furthermore, Y1214 displays more prolonged activation when VEGFR-2 is presented with matrix-bound VEGF compared to soluble (Chen et al., 2010). With this previous knowledge of Y1054/Y1059 and Y1214, we investigated how these specific tyrosine residues affect VEGFR-2 expression, activity, and blood vessel growth in the context of strain-induced angiogenesis. We utilized CRISPR-based VEGFR-2 mutants with non-phosphorylatable residues at 1054 or 1214 positions in immortalized ECs for both vasculogenic and angiogenic studies in 3D microtissue systems.

To characterize how loss of specific phosphorylation sites altered response to ligand stimulation, the mutant lines were tested in standard cell culture with short-term VEGF treatments (**Fig 2B-G)**. For Y1054F cells, overall VEGFR-2 expression was unchanged in comparison to Cas9 control line with or without exogenous VEGF (**Fig 2B,E**). This indicates that there are no significant differences in total receptor expression levels, demonstrating that the receptor is still present at equivalent levels in mutant versus control lines. The Y1054F samples had lower phosphorylation at 1054F/Y1059 compared to Cas9 samples after 15min treatment with VEGF (**Fig 2C,D**). Similar levels of pY1214 were present in Cas9 and Y1054F samples with VEGF stimulation, indicating that this residue can still be activated in the mutant line. On the other hand, the Y1214F samples showed lower levels of total VEGFR-2 compared to Cas9 after stimulation with exogenous VEGF (**Fig 2E**); this is in line with expectations that activated receptors should be internalized and degraded to prevent feedforward signaling (LaValley et al., 2017a). There were no differences in activation at Y1054 after VEGF stimulation between Cas9 and Y1214 samples (**Fig 2F**) but much lower levels of activation at Y1214 as expected (**Fig 2G**). These initial studies suggest that Y1054F cells may experience partial rescue for VEGFR-2 activation and subsequent signaling by the continued presence of Y1059. Notably, our studies used 25ng/mL VEGF, representing almost 10-fold higher levels compared to what CAFs endogenously produce. Therefore, these studies would be expected to elicit higher responses compared to what would occur in co-culture systems. Overall, these studies only focused on static cultures with biochemical stimulation via exogenous VEGF, which does not account for mechanoactivation that can occur via strain stimulation.

After characterizing the effect of ligand treatments on mutant VEGFR-2 lines, we began to investigate strain treatments using the Flexcell system. To understand how Y1054F or Y1214F mutations altered VEGFR-2 mechanoactivation, we treated these lines with combinations of strain and/or exogeneous VEGF for 15min. For this study, total VEGFR-2 was generally stable in the Y1054F line regardless of treatment, and these levels were somewhat elevated compared to Cas9 samples of the same treatment **(Fig 3A**) However, there were no differences observed in total VEGFR-2 between Cas9 controls and Y1214F samples **(Fig 3A,D)**. While these results seem divergent from the static, exogenous VEGF studies, the difference may be explained in the Flexcell substrates being collagen-I coated. The Flexcell plates are also less stiff than standard tissue culture plastic, further highlighting how matrix mechanics and integrin signaling may contribute to regulation of the VEGFR-2 pathway. While not significantly different, the elevated levels of total VEGFR-2 in Y1054F samples compared to Cas9 samples may indicate a potential role of Y1054 in regulating receptor expression. The Y1054F samples show abrogated activation of VEGFR-2 at Y1214 in response to strain and/or VEGF **(Fig 3C)**. Overall, these results both confirmed the loss of Y1214 in the mutant Y1214F line as a functional change and displayed an altered VEGFR-2 expression and activity pattern with the loss of Y1054 or Y1214 in response to VEGF, strain, or the combination of these stimuli. In addition to changes in receptor expression and activation, we investigated the roles of these specific residues in altering blood vessel growth in 3D microtissue models of the TME.

Anti-cancer therapies such as bevacizumab inhibit angiogenesis by blocking the binding of VEGF to VEGFR-2. In our studies, we first looked to evaluate if bevacizumab was sufficient to decrease levels of VEGF secreted by cells embedded in the microtissues, as well as whether it was effective in suppressing vessel growth with mechanical stimulation (**Fig 4A,B**). CAFs secreted high levels of VEGF secretion which was significantly decreased by increasing concentrations of bevacizumab (**Fig 4A**). In modified and unmodified HUVEC lines, there were noticeably lower levels of secreted VEGF, and these levels were not significantly affected at any level of bevacizumab treatment. In 3D microtissue assays, bevacizumab decreased average vessel length in samples with CAFs+HUVEC TERT2s or Cas9 ECs but not in any of the VEGFR-2 mutant samples (**Fig 4B, S3A**). Our lab has previously shown that mechanical and chemical stimulation can alter VEGFR-2 expression and activation, which ultimately alters vasculogenesis (Johnson et al., 2023). To explore biochemical and biomechanical stimuli independently, 3D microtissue assays with HUVEC TERT2 monocultures or HUVEC TERT2 and CAF co-cultures were treated with bevacizumab, exogeneous VEGF, magnetic beads, or combinations of these treatments to determine the effects of on vessel growth (**Fig 4C, S3B,C**). Decreased average vessel length was observed in HUVEC TERT2 monocultures with VEGF plus bevacizumab treatment in comparison to VEGF alone; however total vessel length per area was maintained in samples that received mechanical stimulation from magnetic beads (**Fig 4C, S3C**). These studies highlighted the complexity of blood vessel growth and regulation as a result of combinatorial ligand and/or mechanical cues, potentially suggesting why current anti-angiogenic strategies fail to provide significant improvements in clinical populations despite promising preclinical data.

For vasculogenesis studies performed in fibrin gel disks, NHLFs supported limited blood vessel growth in Y1054F or Y1214F samples **(Fig 5A,B)**. However, Cas9 controls had significantly more vessel growth compared to both mutant lines; this suggests that Y1054 and Y1214 residues are each individually necessary for vascular growth to occur in our models. Y1054/Y1059 are commonly reported to be necessary for full kinase activity of VEGFR-2, and loss of VEGFR-2 has been shown to be embryonically lethal due to insufficient vasculogenesis (Kendall et al., 1999, Shalaby et al., 1995). Therefore, it is unsurprising that Y1054F cells have severely abrogated vascularization potential. However, our results demonstrate that Y1214 is also required for vessel formation. Next, we wanted to replace fibroblasts with ECM microstrains through the addition of magnetic beads to observe how strain would affect these EC lines; these studies were designed to eliminate stromal secreted factors, specifically VEGF from the CAFs or NHLFs, and focus only on mechanical stimulation of vessel behaviors. Both Y1054F and Y1214F lines without magnetically-induced strains had somewhat decreased vessel growth compared to Cas9. Moreover, mechanical stimulation through the magnetic beads was not sufficient to rescue vessel growth in Y1054F or Y1214F samples **(Fig 5C,D)**. Because previous research has shown that strain is sufficient to induce vasculogenesis even without stromal cells, our studies suggest that mechanoactivation of VEGFR-2 at Y1054 or Y1214 may be required for vascular growth (Sewell-Loftin et al., 2017). Our studies are the first to demonstrate that loss of Y1054 or Y1214 reduces the impact of strain on vasculogenesis, highlighting that VEGFR-2 requires both these sites for normal EC function, particularly in response to mechanical strain which is present in the TME due to CAFs.

To take these experiments further, we transitioned from focusing on vasculogenesis to studying angiogenesis through our three-chamber microfluidic devices. Center chambers of devices with NHLFs co-cultured with Cas9 cells showed increased vessel growth compared to chambers with either the Y1054F or Y1214F lines, confirming that vasculogenesis is inhibited by loss of Y1054 or Y1214 **(Fig 6B, E)**. The primary difference between the center chamber and the static ring assays is that the microfluidic models include low levels of interstitial fluid flow (<1µm/s), representing a more physiologically relevant microtissue model (Sewell-Loftin et al., 2020). The Cas9 line also showed the highest levels of angiogenesis into both the NBF and CAF side chambers, with extremely limited angiogenesis for both the Y1054F and Y1214F lines **(Fig 6C)**. There were no significant differences of angiogenesis into NBF chambers compared to CAF chambers for Cas9 samples, suggesting the type of stromal cell does not make a difference in vascularization potential of these ECs. While prior work suggests that CAFs generally support higher levels of vascularization than NBFs, the parental HUVEC TERT2 cells have been modified via CRISPR editing, without the introduction of gRNA for genetic editing. Therefore, it is not necessarily surprising that they behave differently than primary ECs. Additionally, our quantifications only reflect length of vessel growth into side chambers normalized to total growth in the center chamber for that specific device. We did not analyze vessel maturity, lumen structure, or other features that could be differentially regulated by the CAF mechanical stimulation. To further eliminate secreted factors from stromal cells from our studies, we next used the magnetic bead assay to generate micro-strains in the ECM. Interestingly, in the device study with magnetic beads producing matrix strain, Cas9 presented with highest angiogenesis in the chamber with beads and a magnet, emphasizing the pro-angiogenic effects of strain **(Fig 6F)**. However, Y1054F and Y1214F lines continued to show limited angiogenesis, which confirms that loss of these phosphorylation sites is detrimental to EC vessel growth.

Surprisingly, there was some limited rescue of angiogenic potential in both Y1054F and Y1214F samples with mechanical stimulation via magnetic beads. These results are also particularly interesting because interstitial flow in the devices is traveling from the center chamber into the side chamber, and blood vessels are growing in the same direction as the flow. This behavior is different from normal angiogenic activity, where vessels typically travel against the flow gradient (Shirure et al., 2017, Ghaffari et al., 2017). According to these studies, loss of phosphorylation at either tyrosine residue 1054 or 1214 is may partially block angiogenesis via mechanisms related to flow-induced mechanotransduction. The current study expands on this understanding by investigating the roles of Y1054 and Y1214 in strain-induced both angiogenesis and vasculogenesis. Understanding the essential roles of Y1054 and Y1214 in blood vessel growth and tumor progression due to mechanical strain is an important aspect of developing novel anti-angiogenic drugs.

Current anti-angiogenic therapies focus on VEGFR-2 biochemical signaling through the VEGF ligand which is produced by surrounding stromal and tumor cells. However, CAFs present in the TME create strain, which likely mechanically activates VEGFR-2 to promote EC angiogenesis, even in the presence of these anti-angiogenic therapies. This paper confirms the effect that CAF-induced strain has on VEGFR-2 signaling and vessel growth and demonstrates the essential roles of Y1054 and Y1214 in VEGFR-2 activity specifically in response to strain. Strategies such as normalizing TME mechanics by reducing CAF contractility and/or targeting Y1054 and Y1214 could improve the efficacy of modern cancer treatments, allowing for decreased tumor progression and increased survival rate. The techniques and conclusions presented in this paper should be utilized to develop novel and more effective cancer therapies.

## Materials and Methods

### Cell Culture

Primary human umbilical vein endothelial cells (HUVECs) (Lonza, C2517A) and normal human lung fibroblasts (NHLFs) (Lonza, CC-2512) were used between passages 3-6. HUVECs and the immortalized HUVEC TERT2s (ATCC, CRL-4053), along with the HUVEC TERT2 mutant lines utilized (Cas9, Y1054F, Y1214F), were grown in EGM-2 (Lonza, CC-3162). HUVECs and HUVEC TERT2 cells were chosen as the models for studying vasculogenesis and angiogenesis based on substantial use of such models for *in vitro* vessel studies (Nakatsu et al., 2007, Nakatsu et al., 2003, Cao, 2025, Ali et al., 2025, Takano et al., 2008). The use of both primary cells and a derived immortalized line allows for comparison between populations. Previously developed immortalized cancer-associated fibroblasts (CAFs) from a breast cancer patient along with immortalized normal breast fibroblasts (NBFs) from the same patient were used to represent normal and cancerous tissues (Zhang et al., 2016, Corsa et al., 2016). NHLFs, CAFs, and NBFs were grown in DMEM (Gibco, 11995065) supplemented with 1% non-essential amino acid solution (Gibco, 1140-050), 2mM L-glutamine (Gibco, 25-030-081), 100 U/mL penicillin/streptomycin (Fisher Scientific, 15-140-122), 1% sodium pyruvate (Gibco, 11360070), and 10% non-HI FBS (Gibco, 26140079). For cell harvesting, 0.25% trypsin (Gibco 25-200-072) was used. Cells were incubated at 37°C and 5% CO_2_. ECs were treated with exogenous VEGF (Peprotech, 100-20) in cell culture media at 25ng/mL. The myosin II inhibitor blebbistatin (Sigma-Aldrich, 203391) was dissolved in DMSO (Sigma-Aldrich, D8418), and cells were treated with blebbistatin at 20µM, 50µM, or equivalent volumes of DMSO as a vehicle control.

### Mutant Cell Line Clonal Selection

HUVEC TERT2s were modified via CRISPR/Cas9 mutation to generate mutant VEGFR-2 receptor lines (Synthego). For each mutant line, a single nucleotide in the VEGFR-2 coding region was altered to convert either tyrosine 1054 or 1214 to phenylalanine. These mutant lines were labeled as Y1054F and Y1214F, respectively. A control Cas9 line which contained the Cas9 enzyme without any genetic mutations was utilized as a control. To confirm homozygous mutation, we created clonal populations of both Y1054F and Y1214F cells. For each line, 16k cells were seeded in one well of a 96 well plate. A serial dilution was performed throughout the plate, and only wells with single cells and colonies were continuously fed. Once colonies covered at least a third of the well, they were expanded, and DNA was extracted using the QIAwave DNA Blood & Tissue Kit (Qiagen, 69554). Regions surrounding the sites of interest were amplified using the HotStartTaq Master Mix Kit (Qiagen, 203443), and PCR products were purified using QIAquick PCR Purification Kit (Qiagen, 28104). Primers were from IDT **(Table 1)**. Finished PCR samples were sent to the University of Alabama at Birmingham Heflin Center for Genomic Science for Sanger sequencing. Resulting data was analyzed using BioEdit.

**Table 1.**
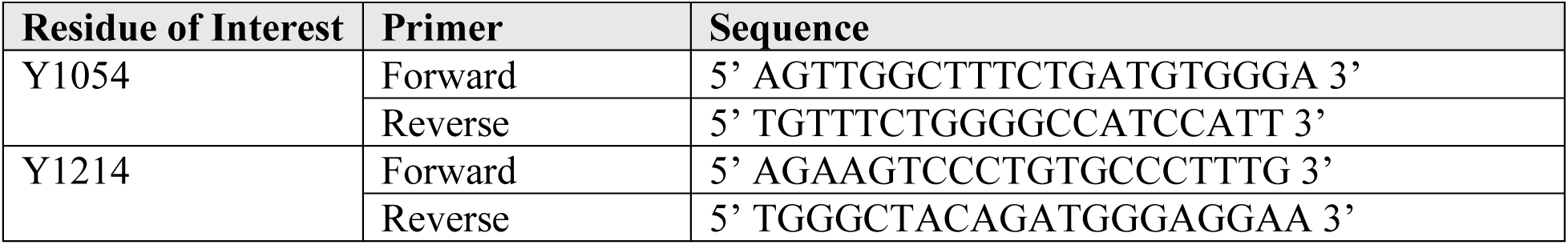
Y1054 and Y1214 Primers. Primer sequences utilized to run PCR and Sanger sequencing on Cas9, Y1054F, and Y1214F lines.
Forward primers were used for Sanger sequencing.

### 3D Fibrin Microtissue Assay

A 3D TME microtissue model was previously developed in our lab with fibrin gel disks formed in a 1cm ring of polydimethylsiloxane (PDMS) (Dow Corning, Sylgard 184) placed on a glass coverslip (Fisher, 22-293232) (Sewell-Loftin et al., 2017). Rings with the coverslips were autoclaved and placed into a 24 well plate before being loaded with fibrin gels embedded with cells. Fibrinogen (Sigma-Aldrich) was solubilized in DPBS without magnesium or calcium ions. For vasculogenesis studies, 50k ECs and 50k fibroblasts were resuspended in a final concentration of 10mg/mL fibrinogen with 3U/mL thrombin before being loaded into a ring. Experiments with magnetic beads included 1.7µL of thrombin-coated magnetic beads per ring. Rings incubated at 37°C before being fed with 1mL EMG-2. Samples were grown for 7d, with media being changed every 2 days. After 7d, rings were fixed for immunofluorescence analysis of vessel growth.

### Microfluidic Devices

Three-chamber microfluidic devices were previously designed to study a variety of cell functions including angiogenesis (Sewell-Loftin et al., 2020). The chambers are in series (now to be referred to as left, center, and right) and fluid lines above and below each chamber allow for control of the direction of media flow between regions. A 10mg/mL fibrin gel is injected into each chamber separately, which may also have cells or magnetic beads embedded in the gel depending on the experiment. For the angiogenesis studies presented in this paper, ECs and NHLFs were seeded into the center at 1.0 x 10^7^cells/mL in a 1:1 ratio. Experiments with CAFs and NBFs in the side chambers were loaded at a concentration of 1.25 x 10^7^cells/mL. Experiments with thrombin-coated magnetic beads and blank fibrin gels were utilized to test angiogenesis as a result of cell-free strain. Some of these devices were grown above a rotating magnet, creating strains in the bead chamber, while others were not and acted as controls. Devices were cultured for 7d at 37°C and 5% CO_2_ then fixed with formalin.

### Matrix Deformation Assay

To analyze CAF contractility, CAFs were loaded into 3D 10mg/mL fibrin gels with 75k cells per gel. Samples were cultured with 1µm blue fluorescent polystyrene microbeads (Invitrogen, F8815). After growing for 24hr, CAFs were treated with full DMEM as described above with no treatment (NT), a DMSO control (Vehicle, Veh), 20µM blebbistatin, or 50µM blebbistatin. After 24hr of treatment, rings were imaged at 30min intervals with an Olympus IX83 microscope as described above using a 100µm Z-stack with 1µm slices. The microscope was equipped with an environmental chamber allowing for incubation at 37°C and 5% CO_2_ to allow for the extended period of imaging. Bead displacement was calculated from the resulting images using a MatLab program, which correlated with matrix deformation caused by the CAFs (Sewell-Loftin et al., 2017).

### Flexcell Strain Studies

A Flexcell system (Flexcell, FX-6000 T) was utilized to apply uniaxial strain on HUVEC TERT2 Cas9, Y1054F, and Y1214F lines. Cells were plated on collagen I coated uniaxial Flexcell plates (Flexcell International Corporation, UF-4001C) at 500k cells per well in EGM-2 48hr before treatment. For the no treatment (NT) group, cell media on the Flexcell plates was changed to EGM-2 without added VEGF (0ng/mL) for 15min. The media for the Strain treatment groups was also changed to EGM-2 with 0ng/mL VEGF, but they were then immediately strained for 15min. To study the interaction of biochemical and biomechanical stimulation VEGF Only and Strain+VEGF treatment groups received a 15min of EGM-2 with 25ng/mL VEGF. Groups treated with Strain were treated with oscillatory strain at 10% elongation and 0.3Hz for 15min, concurrent with either the 0ng/ml or 25ng/mL VEGF treatment. The elongation was chosen to mimic microstrains produced by CAFs in previous studies, and the rate reflects average human respiration (Sewell-Loftin et al., 2017). Samples were run in triplicate.

### Magnetic Microbeads

Iron oxide beads (Dynabeads, M-450 ThermoFisher, 14013) of 5µm diameter and a tosyl-group surface coating were coated by thrombin in preparation to be incorporated into fibrin gels as previously described (Sewell-Loftin et al., 2017). Briefly, beads were coated by washing 100µL of bead stock solution with DPBS, then resuspended in 100µl sterile bicarb buffer (Sigma-Aldrich, C3041) and 100µL 50U/mL thrombin (Sigma-Aldrich, T4648) in DPBS with 0.1% BSA (Sigma-Aldrich, A2153). This mixture was rotated overnight at 4°C, then washed twice in 0.1% BSA in DPBS before being resuspended in 200µL of the 0.1% BSA in DPBS. This method allows for the addition of cell-free mechanical strains in 3D culture systems to ensure stromal cell secretions are not contributing to activation of signaling pathways. Since the beads are effectively crosslinked in the fibrin gel, an external moving magnet will cause bead movement, mimicking matrix deformations caused by CAFs and other stromal cells. In our studies, an orbital plate with a magnet is placed below fibrin gels with the magnetic beads to replicate strain, and control samples are with beads but no magnet.

### Molecular Analyses

For analysis of VEGF secretion by ELISA, 100k CAFs were grown in 3D microtissues for 2d before being treated with a DMSO vehicle control or 20µM blebbistatin in 0ng/ml VEGF EGM-2 for 2d; a no treatment group received only 0ng/ml VEGF EGM-2 for 4d total. This media was harvested. 2D CAFs and HUVECs were also plated along with HUVECs grown in 3D culture gels with 100k or 200k cells per gel. EGM-2 without added VEGF was harvested after two days on the cells. VEGF levels were analyzed using the Human VEGF ELISA Kit (Abcam, ab222510). Plate reader results were normalized to a standard line. In a different VEGF ELISA, HUVEC TERT2s, HUVEC TERT2 Cas9, Y1054F, Y1214F, CAFs, and HUVEC TERT2 and CAF co-cultures were grown in 3D fibrin ring assays for 7d.

Microtissues were grown for 2d before being treated with a DPBS vehicle control (Veh), Bevacizumab (at 2.5µg/mL, 25µg/mL, or 250µg/mL), with media changes every 2d. Conditioned media was harvested on day 6, and VEGF levels were analyzed using the Human VEGF ELISA Kit as described above.

Protein levels were compared using western blot analyses. Lysates were made using RIPA buffer with 50mM Tris pH 7.4, 150mM NaCl (Sigma-Aldrich, S3014), 1% Triton X-100 (Sigma-Aldrich, T8797), 0.25% sodium deoxycholate (Sigma-Aldrich, L3771), 5mM sodium fluoride (Sigma-Aldrich, S6776), and 1mM EDTA (Sigma-Aldrich, E9884) with additional 1:100 HALT protease and phosphate inhibitor (ThermoScientific, 78441). Standard western blot protocols were followed. Protein concentrations were quantified by performing a bicinchoninic acid (BCA) protein assay (ThermoScientific, 23227), and 50µg of protein were loaded per well. Lysates were run on 10% SDS-PAGE gels then transferred onto PDVF membranes. These were blocked in 5% milk in TBST then stained for VEGFR-2 (1:1000, Cell Signaling Technology, 2479), VEGFR-2 pY1054/Y1059 (1:500, Invitrogen, 44-1047; 1:1000, Abcam, ab5473), VEGFR-2 pY1214, and β-actin (1:40,000, Sigma-Aldrich, A1978). Incubations were overnight on a rotator plate at 4°C. Secondary antibodies (Cell Signaling Technology, 7074, 7076) were diluted at 1:2000 in 5% milk in TBST, and membranes were incubated at room temperature for 2 hours on a rotator plate with the secondary. Chemiluminescent detection (ECL, ThermoScientific, 32106; Femto, ThermoScientific, 34095) was used to image the blots on a Licor system, and images were analyzed via densitometry through FIJI. β-actin was used for loading control.

### Immunofluorescence Staining and Imaging

Fibrin gels from both the 3D ring assay and microfluidic devices were stained with CD31 (1:200, Fisher Scientific, PIMA513188) and Alexa Fluor 555 (1:500, ThermoFisher, A31570) to mark ECs forming vasculature. Fixation and staining for rings were performed on a rotator plate, while devices used pressure differences to create flow of solutions. Rings were washed with 1X PBS (Fisher Scientific, BP39920) then fixed with 10% formalin (Fisher Scientific, SF100-4) for 20min at room temperature. They were washed again with 1x PBS and blocked with Abdil (PBS plus 2% BSA and 0.1% Tween-20) for 1hr at room temperature, then they were incubated overnight in CD31 in Abdil at 4°C. After 4 washes, rings were incubated overnight at 4°C in Alexa Fluor 555 in Abdil then washed again 4 times.

At 4°C, microfluidic devices were fixed with 10% formalin for 2d. They were blocked with Abdil for 2d then stained with CD31 in Abdil for 2d. After a wash for 1d, devices were stained with Alexa Fluor 555 in Abdil for 2d then washed. Samples were imaged at 4x for Sytox quantification and 10x for all other studies with an inverted epifluorescence Olympus Microscope (IX83), capturing a 100µm z-stack with a step size of 2µm. Using FIJI, 2D images of microtissues were created using Extended Depth of Field, and max-z projections were created and stitched together for devices to capture all three chambers (Preibisch et al., 2009). Vascular growth was quantified using both FIJI and AngioTool (Zudaire et al., 2011). For vasculogenic rings, total vessel length divided by explant area and average vessel length were calculated. For devices, total vessel length of the side chambers was normalized to total vessel length in the center chamber.

### Statistical Analysis

Data are reported as averages + SEM with 2-9 replicates per group. Graphs show individual data points. To determine statistical significance, data was screened for normality using Shapiro-Wilk tests. If normal, we performed ANOVA with post hoc Tukey HSD tests. If any groups had non-normal sample distribution, we performed Kruskal-Wallis tests with post hoc Dunn’s tests. Statistical calculations were completed using Real Statistics Resource package for Excel. Statistical significance was defined as p < 0.05.

Primer sequences utilized to run PCR and Sanger sequencing on Cas9, Y1054F, and Y1214F lines. Forward primers were used for Sanger sequencing.

## Statements and Declarations

### Data Availability

All data is presented in the main article or in the Additional Files. Datasets will be available upon request to Dr. Sewell-Loftin.

### Author Contributions

Bronte Johnson – Conceptualization, Methodology, Validation, Investigation, Formal Analysis, Writing – Original Draft, Writing – Review & Editing; Tori McKinley – Conceptualization, Methodology, Validation, Investigation, Formal Analysis, Writing – Original Draft, Writing – Review & Editing; Thomas Nguyen – Investigation, Formal Analysis, Writing – Review & Editing; Eden Beasley-Duncan – Investigation, Formal Analysis, Writing – Review & Editing; Tanvi Gridhar – Investigation, Formal Analysis, Writing – Review & Editing; M.K. Sewell-Loftin – Conceptualization, Validation, Formal Analysis, Writing – Original Draft, Writing – Review & Editing, Supervision, Resources, and Funding Acquisition.

## Acknowledgements

Funding Acknowledgements

The authors wish to thank the following funding sources: The American Cancer Society (RSG-24- 1321527-01-MM, M.K.S.L.,) the Breast Cancer Research Foundation of Alabama via the O’Neal Invests Program (M.K.S.L.), and the UAB Graduate School Blazer Fellowship (T.M.). We would also like to thank the Heflin Center for Genomic Sciences which is supported by a University Wide Interdisciplinary Research Center (UWIRC) from the University of Alabama at Birmingham. Finally, we would like to thank the KURE Program (R25 DK115353, PI: Dr. Jennifer Pollock) for supporting E.B.D. and T.G. in summer internships where this work was completed.

## References

Ahmad, A. & Nawaz, M. I. 2022. Molecular mechanism of VEGF and its role in pathological angiogenesis. Journal of Cellular Biochemistry, 123, 1938–1965.

Alcoser, T. A., Bordeleau, F., Carey, S. P., Lampi, M. C., Kowal, D. R., Somasegar, S., Varma, S., Shin, S. J. & Reinhart-King, C. A. 2015. Probing the biophysical properties of primary breast tumor-derived fibroblasts. Cell Mol Bioeng, 8, 76–85.

Ali, A., Roy, B., Schott, M. B. & Grove, B. D. 2025. AKAP12 Variant 1 Knockout Enhances Vascular Endothelial Cell Motility. J Vasc Res, 62, 312–329.

Ayad, N. M. E., Kaushik, S. & Weaver, V. M. 2019. Tissue mechanics, an important regulator of development and disease. Philos Trans R Soc Lond B Biol Sci, 374, 20180215.

Bala, V., Patel, V. & Sewell-Loftin, M. K. 2024. Cadherin Expression Is Regulated by Mechanical Phenotypes of Fibroblasts in the Perivascular Matrix. Cells Tissues Organs, 1–18.

Bates, M. E., Libring, S. & Reinhart-King, C. A. 2023. Forces exerted and transduced by cancer-associated fibroblasts during cancer progression. Biol Cell, 115, e2200104.

Cao, Y. 2025. Human Umbilical Vein Endothelial Cells (HUVECs) in Pharmacology and Toxicology: A Review. J Appl Toxicol, 45, 2512–2545.

Carmeliet, P. & Jain, R. K. 2000. Angiogenesis in cancer and other diseases. Nature, 407, 249–57.

Chen, T. T., Luque, A., Lee, S., Anderson, S. M., Segura, T. & Iruela-Arispe, M. L. 2010. Anchorage of Vegf to the extracellular matrix conveys differential signaling responses to endothelial cells. The Journal of cell biology, 188, 595–609.

Corsa, C. A., Brenot, A., Grither, W. R., Van Hove, S., Loza, A. J., Zhang, K., Ponik, S. M., Liu, Y., Denardo, D. G., Eliceiri, K. W., Keely, P. J. & Longmore, G. D. 2016. The Action of Discoidin Domain Receptor 2 in Basal Tumor Cells and Stromal Cancer-Associated Fibroblasts Is Critical for Breast Cancer Metastasis. Cell Rep, 15, 2510–23.

Dessalles, C. A., Babataheri, A. & Barakat, A. I. 2021. Pericyte mechanics and mechanobiology. J Cell Sci, 134.

Dey, N., De, P. & Brian, L. J. 2015. Evading anti-angiogenic therapy: resistance to anti-angiogenic therapy in solid tumors. Am J Transl Res, 7, 1675–98.

Ewan, L. C., Jopling, H. M., Jia, H., Mittar, S., Bagherzadeh, A., Howell, G. J., Walker, J. H., Zachary, I. C. & Ponnambalam, S. 2006. Intrinsic tyrosine kinase activity is required for vascular endothelial growth factor receptor 2 ubiquitination, sorting and degradation in endothelial cells. Traffic, 7, 1270–82.

Folberg, R., Hendrix, M. J. & Maniotis, A. J. 2000. Vasculogenic mimicry and tumor angiogenesis. Am J Pathol, 156, 361–81.

Ghaffari, S., Leask, R. L. & Jones, E. A. V. 2017. Blood flow can signal during angiogenesis not only through mechanotransduction, but also by affecting growth factor distribution. Angiogenesis, 20, 373–384.

Hashimoto, T. & Shibasaki, F. 2015. Hypoxia-Inducible Factor as an Angiogenic Master Switch. *Frontiers in Pediatrics*, Volume 3–2015.

Johnson, B. M., Johnson, A. M., Heim, M., Buckley, M., Mortimer, B., Berry, J. L. & Sewell- Loftin, M. K. 2023. Biomechanical stimulation promotes blood vessel growth despite Vegfr-2 inhibition. Bmc Biol, 21, 290.

Kendall, R. L., Rutledge, R. Z., Mao, X., Tebben, A. J., Hungate, R. W. & Thomas, K. A. 1999. Vascular endothelial growth factor receptor Kdr tyrosine kinase activity is increased by autophosphorylation of two activation loop tyrosine residues. J Biol Chem, 274, 6453–60.

Koch, M. K., Jaeschke, A., Murekatete, B., Ravichandran, A., Tsurkan, M., Werner, C., Soon, P., Hutmacher, D. W., Haupt, L. M. & Bray, L. J. 2020. Stromal fibroblasts regulate microvascular-like network architecture in a bioengineered breast tumour angiogenesis model. Acta Biomater, 114, 256–269.

Koch, S. & Claesson-Welsh, L. 2012. Signal transduction by vascular endothelial growth factor receptors. Cold Spring Harb Perspect Med, 2, a006502.

Labrecque, L., Royal, I., Surprenant, D. S., Patterson, C., Gingras, D. & Beliveau, R. 2003. Regulation of vascular endothelial growth factor receptor-2 activity by caveolin-1 and plasma membrane cholesterol. Mol Biol Cell, 14, 334–47.

Lamalice, L., Houle, F., Jourdan, G. & Huot, J. 2004. Phosphorylation of tyrosine 1214 on VEGFR2 is required for Vegf-induced activation of Cdc42 upstream of SAPK2/p38. Oncogene, 23, 434–45.

Lavalley, D. J., Zanotelli, M. R., Bordeleau, F., Wang, W., Schwager, S. C. & Reinhart-King, C. A. 2017a. Matrix Stiffness Enhances Vegfr-2 Internalization, Signaling, and Proliferation in Endothelial Cells. Converg Sci Phys Oncol, 3, 044001.

Lavalley, D. J., Zanotelli, M. R., Bordeleau, F., Wang, W., Schwager, S. C. & Reinhart-King, C. A. 2017b. Matrix Stiffness Enhances Vegfr-2 Internalization, Signaling, and Proliferation in Endothelial Cells. Converg Sci Phys Oncol, 3.

Lugano, R., Ramachandran, M. & Dimberg, A. 2020. Tumor angiogenesis: causes, consequences, challenges and opportunities. Cell Mol Life Sci, 77, 1745–1770.

Miller, B. & Sewell-Loftin, M. K. 2022. Mechanoregulation of Vascular Endothelial Growth Factor Receptor 2 in Angiogenesis. *Frontiers in Cardiovascular Medicine*, Volume 8–2021.

Nakatsu, M. N., Davis, J. & Hughes, C. C. 2007. Optimized fibrin gel bead assay for the study of angiogenesis. J Vis Exp, 186.

Nakatsu, M. N., Sainson, R. C., Aoto, J. N., Taylor, K. L., Aitkenhead, M., Perez-Del-Pulgar, S., Carpenter, P. M. & Hughes, C. C. 2003. Angiogenic sprouting and capillary lumen formation modeled by human umbilical vein endothelial cells (Huvec) in fibrin gels: the role of fibroblasts and Angiopoietin-1. Microvasc Res, 66, 102–12.

Preibisch, S., Saalfeld, S. & Tomancak, P. 2009. Globally optimal stitching of tiled 3d microscopic image acquisitions. Bioinformatics, 25, 1463–5.

Ribatti, D. 2010. The inefficacy of antiangiogenic therapies. J Angiogenes Res, 2, 27.

Ribatti, D. 2016. Tumor refractoriness to anti-Vegf therapy. Oncotarget, 7, 46668–46677.

Roman, B. L. & Pekkan, K. 2012. Mechanotransduction in embryonic vascular development. Biomech Model Mechanobiol, 11, 1149–68.

Roskoski, R., Jr. 2007. Vascular endothelial growth factor (Vegf) signaling in tumor progression. Crit Rev Oncol Hematol, 62, 179–213.

Sasich, L. D. & Sukkari, S. R. 2012. The Us FDAs withdrawal of the breast cancer indication for Avastin (bevacizumab). Saudi Pharm J, 20, 381–5.

Sewell-Loftin, M. K., Bayer, S. V. H., Crist, E., Hughes, T., Joison, S. M., Longmore, G. D. & George, S. C. 2017. Cancer-associated fibroblasts support vascular growth through mechanical force. Sci Rep, 7, 12574.

Sewell-Loftin, M. K., Katz, J. B., George, S. C. & Longmore, G. D. 2020. Micro-strains in the extracellular matrix induce angiogenesis. Lab Chip, 20, 2776–2787.

Shalaby, F., Rossant, J., Yamaguchi, T. P., Gertsenstein, M., Wu, X. F., Breitman, M. L. & Schuh, A. C. 1995. Failure of blood-island formation and vasculogenesis in Flk-1-deficient mice. Nature, 376, 62–6.

Shirure, V. S., Lezia, A., Tao, A., Alonzo, L. F. & George, S. C. 2017. Low levels of physiological interstitial flow eliminate morphogen gradients and guide angiogenesis. Angiogenesis, 20, 493–504.

Simons, M. 2012. An inside view: Vegf receptor trafficking and signaling. Physiology (Bethesda), 27, 213–22.

Takano, H., Murasawa, S. & Asahara, T. 2008. Functional and gene expression analysis of htert overexpressed endothelial cells. Biologics, 2, 547–54.

Testini, C., Smith, R. O., Jin, Y., Martinsson, P., Sun, Y., Hedlund, M., Sainz-Jaspeado, M., Shibuya, M., Hellstrom, M. & Claesson-Welsh, L. 2019. Myc-dependent endothelial proliferation is controlled by phosphotyrosine 1212 in Vegf receptor-2. Embo Rep, 20, e47845.

Van Beijnum, J. R., Nowak-Sliwinska, P., Huijbers, E. J., Thijssen, V. L. & Griffioen, A. W. 2015. The great escape; the hallmarks of resistance to antiangiogenic therapy. Pharmacol Rev, 67, 441–61.

Vion, A. C., Perovic, T., Petit, C., Hollfinger, I., Bartels-Klein, E., Frampton, E., Gordon, E., Claesson-Welsh, L. & Gerhardt, H. 2020. Endothelial Cell Orientation and Polarity Are Controlled by Shear Stress and Vegf Through Distinct Signaling Pathways. Front Physiol, 11, 623769.

Yu, H., Mouw, J. K. & Weaver, V. M. 2011. Forcing form and function: biomechanical regulation of tumor evolution. Trends Cell Biol, 21, 47–56.

Zanotelli, M. R. & Reinhart-King, C. A. 2018. Mechanical Forces in Tumor Angiogenesis. Adv Exp Med Biol, 1092, 91–112.

Zhang, K., Grither, W. R., Van Hove, S., Biswas, H., Ponik, S. M., Eliceiri, K. W., Keely, P. J. & Longmore, G. D. 2016. Mechanical signals regulate and activate SNAIL1 protein to control the fibrogenic response of cancer-associated fibroblasts. J Cell Sci, 129, 1989–2002.

Zudaire, E., Gambardella, L., Kurcz, C. & Vermeren, S. 2011. A computational tool for quantitative analysis of vascular networks. PLos One, 6, e27385.

